# Antibody Transcytosis and Neutralizing Activity in Respiratory Epithelial Cells

**DOI:** 10.64898/2026.05.25.727697

**Authors:** Eduardo U. Anaya, Yee Vue, Maggie Li, Jessica D. Resnick, Nico J. Swanson, David J. Sullivan, Andrew Pekosz

## Abstract

The neutralizing activity present in human serum is considered a correlate of protection against SARS-CoV-2 infection and disease but the mechanisms by which serum antibodies are transported into the lumen of the respiratory tract, where they are required to interact with virus particles and infected cells remain incompletely understood. The transcytosis and neutralizing activity of serum-derived IgG and IgA antibodies was investigated using an in vitro SARS-CoV-2 infection model with primary differentiated human nasal and basal epithelial cells (hNECs and hBECs) cultures. Expression of the antibody transport receptors neonatal Fc receptor (FcRn) and polymeric immunoglobulin receptor (pIgR) in hNECs cultures was confirmed by qPCR, immunofluorescence microscopy, and flow cytometry. Both receptors were expressed throughout the epithelial cultures, with enriched expression observed in ciliated cells compared with goblet and basal cells. Purified IgG and IgA isolated from convalescent plasma demonstrated specificity for SARS-CoV-2 spike protein and inhibited ACE2-Spike interactions, although activity was reduced against later variants. Purified IgG contained higher anti-spike antibody titers than purified IgA. Functional neutralization assays showed that transcytosed IgG and IgA significantly reduced SARS-CoV-2 infection compared with untreated controls. However, serial dilution studies demonstrated that IgG-mediated neutralization was more potent than IgA-mediated neutralization. Similar results were determined for influenza A virus H3N2 subtype. The transcytosis of IgG was more efficient in hBEC cultures while IgA transcytosis was higher in hNEC cultures, reflecting the levels of the corresponding transport proteins. Together, these findings demonstrate that serum-derived IgG and IgA can undergo transepithelial transport across human nasal epithelium while retaining SARS-CoV-2 or influenza A virus neutralizing activity in vitro. These results suggest that FcRn- and pIgR-mediated antibody transport may contribute to mucosal protection following vaccination or infection and may help identify antibody responses associated with protection against SARS-CoV-2.

**Importance:** Serum antibody levels are considered correlates of protection for SARS-CoV-2 and Influenza A virus but its unclear how those antibodies are transported to the apical surface of respiratory epithelial cells, where they must be present for optimal activity. We show that IgG and IgA specific for SARS-CoV-2 or influenza A virus is transcytosed across respiratory epithelial cell cultures, the efficiency of which reflects the level of FcRn or pIgR expression levels, suggesting that cells of the upper and lower respiratory tract transport different antibodies from the blood.

## Introduction

The severe acute respiratory syndrome coronavirus 2 (SARS-CoV-2) is a highly transmissible and pathogenic coronavirus that caused the 2019 coronavirus disease 2019 (COVID-19) pandemic. The sinonasal airway epithelium represents one of the initial sites of SARS-CoV-2 infection and viral replication. The respiratory epithelium consists of a heterogeneous population of ciliated, mucus-secreting goblet, and basal cells that collectively mediate mucociliary clearance^(1)^. Previous studies from our laboratory and others have demonstrated that ciliated and goblet cells are primary cellular targets of SARS-CoV-2 infection and support early viral replication ^(2)^. The viral entry of SARS-CoV-2 begins through host cell receptors, angiotensin-converting enzyme 2 (ACE2), and host serine proteases, including transmembrane serine protease 2 (TMPRSS2). As the primary cells of infection, both nasal and bronchial tissues express ACE2 and TMPRSS2 proteins ^(4)^.

At the initial site of infection, nasal epithelial cells have developed an array of defense mechanisms to protect against invading pathogens. These mucosal defenses consist of both epithelial and immunological barrier components. Previous studies have demonstrated that mucosal epithelia have well-developed tight junctions that contribute to barrier integrity ^(5–8)^. Collectively, the epithelial barrier and mucosal layers prevent invasion through tight junctions and the separation of the host interior from the invading viral pathogens ^(9)^. Additionally, immunological barriers further enhance protection through immunoglobulin antibodies, which prevent binding of the virus to the epithelial cells, suppress viral replication, and reduce virulence ^(10–12)^.

During the early stages of respiratory viral infection, mechanisms that block infection and limit viral replication spread are important for limiting virus spread. Vaccine- and infection-induced neutralizing antibody activity in human serum is considered one of the strongest correlates of protection against SARS-CoV-2 infection and disease ^(13,14)^. Anti-SARS-CoV-2 spike Immunoglobulin G (IgG) and A (IgA) neutralizing antibodies that target the SARS-CoV-2 spike protein confer protection in humans and animals ^(15–22)^. While it is known that SARS-CoV-2 vaccination does not induce a strong mucosal IgA or IgG response in humans, it remains incompletely understood how serum antibodies eventually access the lumen of the upper respiratory tract, where mucosal protection against infection is required.

Within the mucosal tissues, the neonatal Fc receptor (FcRn) and Poly Immunoglobulin Receptor (pIgR) are key receptors that mediate the transportation of immunoglobulins. The FcRn is a major histocompatibility complex (MHC) Class I-related receptor that is semi-ubiquitously expressed in diverse tissues throughout the body, including epithelia, endothelia, and cells of hematopoietic origin^(11,23–25)^. FcRn is responsible for binding IgG and albumin and translocating across polarized endothelium and epithelium cell layers ^(24,25)^. In addition, FcRn protects IgG from lysosomal degradation by directing internalized antibodies into endosomal recycling pathways ^(26)^.

In contrast, pIgR is primarily expressed on epithelial cells of the respiratory and gastrointestinal tract, as well as glandular epithelial tissues, including the breast and liver ^(27,28)^. The receptor functions primarily in trancytosing dimeric IgA and IgM from the basal lamina to the lumen of mucosal tissues ^(27–29)^. On the cell surface, pIgR binds to polymeric IgA and IgM via their J-chain. Ig-pIgR complexes are internalized via clathrin-mediated endocytosis. Internalized complexes go through endosomal pathways that start at early endosomes, then are sorted into common endosomes, and are then rescued to apical recycling endosomes ^(30,31)^.

In this study, the transcytosis and neutralizing activity of serum-derived IgG and IgA antibodies were investigated using primary differentiated human nasal epithelial cell (hNECs) or bronchial epithelial cell (hBEC) cultures infected with SARS-CoV-2 or influenza A virus. Both receptors were detected in differentiated hNEC and hBEC cultures and both epithelial culture systems demonstrated the capacity to transcytose antibodies and inhibit viral infections. Collectively, these findings support the role of FcRn and pIgR in mediating IgG and IgA transcytosis and highlight the utility of primary respiratory epithelial cultures for investigating antibody-mediated neutralization of respiratory viral infections.

## Methods

### Cell Culture

VeroE6/TMPRSS2 cells (VT; RRID: CVCL_YQ49) were obtained from the cell repository of the National Institute of Infectious Diseases, Japan, and are described ^(32)^. TMRPSS2-overexpressing Vero E6 cells were cultured in Dulbecco’s Modified Eagle Medium (DMEM)( Gibco Life Technologies, Cat#11965118) with 10% fetal bovine serum (FBS)(Gibco Life Technologies, Cat# A5256701), 100U penicillin/mL with 100 µg streptomycin/mL (Quality Biological, Cat# 120-095-721), 2mM L-Glutamine (Gibco Life Technologies, Cat# 25030081), and 1m M Sodium Pyruvate (Sigma, Cat# S8636) at 37°C with air supplemented with 5% CO_2_. Infectious medium for SCV2 (IM-SCV2) was used in all infections and consists of DMEM with 2.5% FBS, 100U penicillin/mL with 100 µg streptomycin/mL, 2m M L-Glutamine, and 1mM Sodium Pyruvate.

Madin-Darby Canine Kidney (MDCK) cells were cultured in DMEM containing 10% FBS, penicillin 100 U/ml, streptomycin 100 μg/ml, and 1 mM sodium pyruvate (at 37°C) in a humidified environment with air supplemented with 5% CO_2_. Infection media for IAV (IM-IAV) was used in all infections and consists of DMEM with 4 µg/mL N-acetyl trypsin (NAT)(Sigma, Cat# T6763), 100 ug/mL penicillin with 100 ug/mL streptomycin, 2 mM L-Glutamine, and 0.5% bovine serum albumin (BSA) (Sigma, Cat #A9647).

Human nasal epithelial cells (hNECs) or bronchial epithelial cells (hBECs) (Promocell) were grown to confluence in 24-well Falcon filter inserts (0.4-uM pore; 0.33cm^2^; Becton Dickinson) using PneumaCult™-Ex Plus Medium (Stemcell, Cat# 05001). Confluence was determined by a TEER reading above 250Ω ^(33)^. The cells were then differentiated at an air-liquid interface (ALI) before infection, using ALI medium as basolateral media as previously described ^(34)^.

### RNA Extraction and qPCR

RNA was extracted from hNECs using Trizol (Invitrogen, Cat# 15596026) and the PureLink RNA Mini Kit with on-column DNase treatment (Invitrogen, Cat# 12183018A) according to the manufacturer’s protocol. RNA was then converted to cDNA using the High-Capacity cDNA Reverse Transcription Kit (ThermoFisher, Cat# 4368814) according to the manufacturer’s protocol. The cDNA was diluted to 1:10. qPCR was then run using TaqMan reagents according to the manufacturer’s protocol (Master Mix: Applied Biosystems, Cat# 4369016). Probes used were as follows: PIGR (Applied Biosciences, Cat# hs00922561_m1), FCGRT (Applied Biosciences, Cat# hs00175415_m1), and GAPDH (Applied Biosystems, Cat# hs02786624_g1).

### Viruses

VeroE6/TMPRSS2 cells were used to grow and titrate SARS-CoV-2 variants, as previously described ^(90)^. The virus used in this study is an ancestral B.1 lineage (hCoV-19/USA/DC-HP00007/2020, EPI_ISL_434688), Gamma P.1.17 lineage (hCoV-19/USA/MD-HP03867/2021, EPI_ISL_1468644), Omicron B.1.529 lineage (hCoV-19/USA/MD-HP20874/2021, EPI_ISL_7160424), and an influenza H3N2 vaccine strain (A/Kansas/14/2017 X-327). The SARS-CoV-2 variants were isolated at Johns Hopkins University, and for working stock, VeroE6/TMPRSS2 cells in a T75 or T150 flask were infected with an inoculum at an MOI of 0.001, using a SARS-CoV-2 variant diluted in IM-SCV2. The viral incubation was for one hour at 33°C, and then the viral inoculum was removed, followed by PBS wash. IM was added to the flask, and the flasks were observed over the next couple of days for cytopathic effect (CPE) (approx. 75%). IM supernatant was harvested, clarified by centrifugation at 400g for 10 min, aliquoted, and stored at −65°C. Virus stocks were sequenced and verified by extracting vRNA as described above and detected using quantitative RT-PCR. The consensus sequences of the seed stock and working stock did not differ from the sequence derived from the clinical isolate.

The influenza A vaccine strain working stocks were grown on MDCK cells in a T75 or T150 flask and then infected for one hour at an MOI of 0.001 with the virus diluted in IM-IAV. The flask was observed for CPE (approx. 50%), the supernatant was harvested, centrifuged at 400g for 10 min, aliquoted, and stored at −65°C. Working stocks and vaccine strain were sequenced, and consensus sequences were the same.

### Immunofluorescence Microscopy

Both cell cultures were subjected to immunofluorescence staining as previously described ^(34)^. The wells were washed with PBS and fixed using 4% paraformaldehyde, then permeabilized and blocked with PBS containing 0.5% Triton X-100 and 5% BSA. The wells were immunostained using anti-FcRn (Novus Bio, Cat# NBP1-59061) or anti-pIgR (ThermoFisher, Cat# PA5-35340), and anti-β-Tubulin IV (Abcam, Cat# ab21057) primary antibodies at a 1:100 dilution. Fluorescently labeled secondary antibodies AF488 (ThermoFisher, Cat# A11013) and AF647 (ThermoFisher, Cat# A21244) were used as secondary stains (1:500). The slides were mounted on microscope slides using ProLong^TM^ Glass Antifade Mountant (ThermoFisher, Cat# P36982). Sample images were acquired with a Zeiss LSM700 at 63x magnification with 1 µm z-stack sections. The percentage of FcRn/pIgR cells was quantified using ImageJ ^(35)^.

VeroE6/TMPRSS2 were grown on 8 chamber culture slides and infected with SARS-CoV-2 variants at an MOI of 1. The cells were fixed 24 hr post-infection using 4% paraformaldehyde and permeabilized and blocked with PBS containing 0.5% Triton X-100 and 5% BSA. The cells were stained using purified IgG/IgA, a primary antibody against Spike protein (SinoBiological, Cat# 40590-D001), and Hoechst stain. Fluorescently labeled secondary antibodies AF488 and AF555 were used as a secondary stain. The slides were mounted on microscope slides using ProLong^TM^ Glass Antifade Mountant. Images were acquired using a brightfield Zeiss Axio Imager microscope at 40x magnification and analyzed with ImageJ ^(35)^.

For colocalization analysis, hNECs cultures were harvested from the apical membrane into a single cell suspension by incubating for 30 min in 1X TrypLE (Gibco, Cat# 12563011). TrypLE was then inactivated using stop solution (10% FBS in PBS). Cells were pelleted by centrifugation and resuspended in 1 mL of PBS. A cell suspension of 750,000 cells was made, and 100 µL of suspension was added to the cytospin funnel. Funnel-slide complex was spun at 600 rpm for 4 min. Cells were then fixed with 4% paraformaldehyde for immunofluorescence imaging. The hNECs were stained using anti-FcRn or anti-pIgR, and anti-human IgG (Abcam, Cat# ab200699) or anti-human IgA (ThermoFisher, Cat# SA1-19258) primary antibodies at a 1:100 dilution. To quantify colocalization, Pierce correlation and Mander’s coefficients were determined using the JACoP ImageJ plugin ^(36)^. Pierce correlation is used to describe the correlation of the intensity distributions between two channels ^37^. Mander’s coefficients describe the amount of fluorescence of the colocalizing pixels in each color channel ^(37)^. Pierce and Mander’s correlation coefficients were obtained using Coste’s thresholding ^(38)^.

### Immunoblotting

Low multiplicity of infection (MOI) infections were performed in hNEC cultures. Cells were harvested using TrypLE, lysed, and samples were resolved by SDS-PAGE before transfer onto nitrocellulose membranes. Membranes were probed with primary antibodies against FcRn, pIgR, and β-tubulin. Secondary detection was performed using Alexa Fluor 647- or Alexa Fluor 455-conjugated secondary antibodies or HRP-conjugated secondary antibodies.

### Flow Cytometry

Fully differentiated hNEC or hBEC cultures were harvested from the apical membrane into a single-cell suspension by incubating for 30 min in 1X TrypLE (Gibco, Cat# 12563011). 1X TrypLE was then inactivated using stop solution (10% FBS in PBS). The cells were then washed three times in PBS and resuspended in 1 mL PBS (centrifuged at 600 g for 3 min between wash steps). Appropriate control and sample tubes were then stained with AQUA viability dye (Invitrogen, Cat# L34965) 1 µL/1×10^6^ cells for 30 min at room temperature. Surface staining using anti-FcRn (1:100) or anti-pIgR (1:100) antibodies for 30 minutes on ice.

Fluorescently labeled secondary antibodies AF488 (1:100) were used as a secondary stain and incubated for one hour on ice. Cells were washed, then resuspended in BD Fixation/Permeabilization solution (BD Biosciences, Cat# 554714) and incubated for 30 min at 4°C. Cells were washed with BD Perm/Wash Buffer x2 at 2500 RPM at 4°C for 5 min. Cells were then resuspended in BD Perm/Wash Buffer with 7% Normal Goat Serum (Sigma Aldrich, Cat# G9023) and incubated for 1 hour at 4°C. Cells were washed with BD Perm/Wash Buffer x2 at 600 g at 4°C for 5 min. Appropriate sample tubes were incubated with the primary antibody Anti-Muc5AC (ThermoFisher, Cat# MA5-12178) for one hour at room temperature. Antibodies were diluted into BD Perm/Wash buffer at appropriate concentrations. Final staining volume was 200 µL. Cells were washed with BD Perm/Wash Buffer x2 at 2500 RPM at 4°C for 5 min.

Appropriate sample tubes were incubated with secondary antibodies BV-605 goat anti-mouse (BD Biosciences, Cat# 405327) for 30 min at room temperature. Cells were washed with BD Perm/Wash Buffer x2 at 2500 RPM at 4°C for 5 min. Appropriate sample tubes were incubated with conjugated antibodies, anti-Acetylated alpha tubulin 647 (Santa Cruz, Cat# sc-23950) and Anti-CD271 PE (ThermoFisher, Cat# 12-9400-42) for 30 min at room temperature. Cells were washed with BD Perm/Wash Buffer x2 at 2500 RPM at 4°C for 5 min. Cells were resuspended in FACS Buffer (0.3% BSA in 1X PBS) and filtered through a 35 µm strainer cap into FACS tubes. Cell suspensions were run on a BD LSRII Flow Cytometer using DIVA software. Single-stained cells were used as controls, and a fluorescence minus one control was used to assist in gating. Data analysis was completed on FlowJo V10. Gating strategy was as follows: exclusion of debris, single cells, Aqua-cells, SARS-CoV-2 AF647-cells, and finally BT-IV expression as negative, low, or high (Supplemental Figure 1).

### Antibody Purification

Protein G (InvivoGen, Cat# gel-agg-5) or Peptide M (InvivoGen, Cat# gel-pdm-5) was packed into an appropriate chromatography column to purify either IgG or IgA, respectively. Briefly, 2 mL of the convalescent serum sample was loaded into the column. The columns were washed with buffers containing 10 mM sodium phosphate and 150 mM sodium chloride. The bound antibodies were eluted using a buffer containing 100 mM glycine and then immediately neutralized using a buffer of 1 M TRIS. Purified antibodies were confirmed by Western blot analysis (Supplemental Figure 6).

### Transfection

VeroE6/TMPRSS2 cells were plated to 90% confluency on the day of transfection. A solution of 250 µL of Opti-MEM^TM^ (ThermoFisher, Cat# 31985070, 48 µL of warm TransIT-LT1 (mirusbio, Cat# 2306), and 2 µg of DNA was made. The complexes were incubated at room temperature for 15-30 min. Fresh medium (2% FBS) was added to Vero6/TMPRSS2 cells, and the transfection master mix was added in a drop-wise fashion in different areas of the well. The cells were incubated at 37°C/5% CO2 for 48 hr. After 48 hr post-transfection, cells were washed with PBS and fixed in 4% paraformaldehyde for immunofluorescence imaging.

### Magpix

For these assays, a multilevel “sandwich” was created in duplicate for each sample tested (n = 2). For our 15-plex assay, 15 different types of microspheres were coated with different analytes, which include: Spike proteins from SARS-CoV-2 variants (B.1.1.7, B.1.351, B.1.617.2, P.1), SARS-CoV-2 vaccine components (RBD, Spike trimer, and S1 protein), SARS-CoV2 nucleocapsid, Spike proteins from Seasonal SARS-CoV (229E, HKU1, OC43, NL63, S1) and MERS S1. Briefly, the sandwich consisted of (in order): bead coated in analyte, purified IgG or IgA, detection anti-human Ig conjugated with PE. The coated microspheres were added to 96-well plates. Next, dilutions of purified IgG or IgA were added to the coated microspheres.

The plates were incubated at room temperature (RT) for 2 hr with shaking and then washed two times with the wash buffer. Detection antibodies were then added to the wells for 30 minutes with shaking and then washed two times with the wash buffer. 120 µl of wash buffer was added to each well, shaken for 5 min, and then read.

For the 6-plex ACE2 inhibition assay, 6 different types of microspheres were coated with different analytes which include: Spike proteins from SARS-CoV-2 variants (B.1.1.7, B.1.351, B.1.617.2, P.1, B.1.1.52, and S1). The sandwich consisted of (in order): bead coated in analyte, purified IgG or IgA, ACE2 conjugated with Biotin, detection streptavidin conjugated with PE. The coated microspheres were added to 96-well plates. Next, serial dilutions (0.061-1000 µg/ml) of purified IgG or IgA were added to the coated microspheres. The plates were incubated at room temperature (RT) for 2 hr with shaking, and then washed two times with the wash buffer. ACE2-Biotin was then added to the wells for 30 min with shaking and then washed two times with wash buffer. Detection streptavidin-PE was added for 30 min with shaking and then washed two times with wash buffer. 120 µl of wash buffer was added to each well, shaken for 5 min, then read. The fluorescent signal generated by each well was measured with the Luminex MagPix system. For a result to be valid, a minimum of 50 beads/well had to be detected by the system. IC_50_ values were determined using GraphPad Prism (Supplemental Figure 3).

### Neutralization Assay

Neutralization assays were performed using VeroE6/TMPRSS2 cells for SARS-CoV-2 or MDCK for IAV as previously described ^39,40^. Briefly, antibodies were serially diluted from 1 mg/ml - 3.9 µg/ml at a 1:1 dilution. Samples were then incubated for an hour at room temperature with virus added to plasma at a final concentration of 1 × 10^4^ TCID50/ml. After an hour, 100 μl of each dilution was added to 1 well of a 96-well plate of cells in sextuplet and incubated for 6 hr at 37°C. The inoculum was removed, fresh IM was added, and the plates were incubated at 37°C for 2-4 days until a CPE was visible in the no serum controls. The cells were fixed with 4% formaldehyde, incubated for at least 4 hr at room temperature, and then stained with Naphthol Blue Black (MilliporeSigma, Cat# 195243). As previously described, the nAb titer was calculated as the highest serum dilution that eliminated the CPE in 50% of the wells (NT50) ^(40)^.

### Transepithelial Electrical Resistance (TEER)

Transepithelial electrical resistance (TEER) was measured by using a Millicell ERS-2 Epithelial Volt-Ohm Meter (Millipore Sigma). Briefly, ALI media was removed from the basal compartment of each well, and 150 µL of PBS was added to the apical and basal chambers of the cultures to form an electrical circuit across the cell monolayer and into the basal chamber. Only cultures on trans-wells displaying baseline TEER readings greater than 400 Ω/cm^2^ were used for the experiments ^(33)^.

### FITC Dextran

Paracellular permeability was studied by measuring the apical-to-basolateral flux of FITC- dextran 4 kDa (Sigma, Cat# 46944). Primary differentiated cells that showed TEER readings greater than 400 Ω/cm^2^, the upper chambers were filled with 3 mg/mL of FITC-dextran and incubated for 2 hr at 37 °C. Basolateral media (100 µL) were recovered, serially diluted in a 96-well plate, and fluorescence was measured with a microplate fluorometer at 525 nm (SpectraMax i3x, Molecular Devices, San Jose, CA).

### ELISA

For antibody transcytosis assays, 1 mg/mL of IgG or IgA was added to the basolateral compartment of the hNECs trans well. At each indicated time point, 100 µL of PBS was added to the apical compartment and incubated for 10 min. The concentration of antibodies transcytosed to the apical compartment was quantified using a Human IgG ELISA kit (Abcam; Cat#195215) or Human IgA ELISA kit (Abcam; Cat#196263). Briefly, 10 µL of apical wash sample was diluted in 90 µL of ddH_2_O. Subsequently, 50 µL of diluted sample and 50 µL of detection antibody cocktail were added to a 96-well plate and incubated for 1 h at room temperature with shaking. Plates were then washed 3X with 1X PBS. Next, 100 µL of TMB substrate (Thermo Fisher Scientific; Cat#BDB555214) was added and incubated in the dark for 5 min, followed by the addition of 100 µL stop solution for 1 min. Absorbance was measured at 450 nm using a SpectraMax i3x microplate reader. IC_50_ values were calculated using GraphPad Prism.

### Viral Growth Curves

Prior to hNEC or hBEC infection, the apical side of the transwell was washed with PBS and incubated for 10 min at 37°C. The virus inoculum was diluted in 100 µL of IM at an MOI of 0.25 and added to the apical compartment of hNECs transwell. Following incubation for 2 hr at 37°C, the inoculum was then removed, the cells washed 3x with PBS, and placed in the incubator at 37°C. At each time point, a 10-min apical wash with IM was collected and stored at −80°C. Infectious virus particle production in apical washes was quantified using TCID_50_ on VeroE6/TMPRSS2 or MDCK cells. The basolateral media was collected at the end of the experiment, stored at −80°C.

### TCID_50_ Assay

SARS-CoV-2 infectious virus titers were determined using VeroE6/TMPRSS2 cells, and infectious IAV titers were determined using MDCK-SIAT cells. Cells were seeded into 96-well plates and cultured to 90-100% confluency. After being washed twice with PBS, ten-fold serial dilutions of the viruses in IM-SCV2 or IM-IAV were made, and 20 µL of each dilution was added to its respective cell plates, and each virus dilution was added to the cell plates in six replicates. The plates were incubated at 37°C with 5% CO_2_ for 5-6 days. The cells were fixed by adding 75 µL of 4% formaldehyde in PBS per well overnight and then stained with Napthol Blue Black solution. Endpoint titers were calculated by using the Reed-Muench method ^(41)^.

## Results

### Expression of FcRn and pIgR in primary differentiated nasal epithelial cells

To investigate how FcRn- and pIgR-mediated transport of IgG and IgA across the epithelial cells to neutralization of viral pathogens, we used primary differentiated human nasal epithelial cells (hNECs), a physiologically relevant culture system ^(34)^. Previous studies have reported the expression of FcRn and pIgR in human nasal epithelium ^(44,49)^, and the expression of both receptors was confirmed by quantitative reverse transcription polymerase chain reaction (qRT-PCR) (Figure 1A). Receptor expression was also confirmed by immunofluorescence microscopy using anti-FcRn and anti-pIgR antibodies in hNECs cultures (Figure 1B).

**Figure 1.**
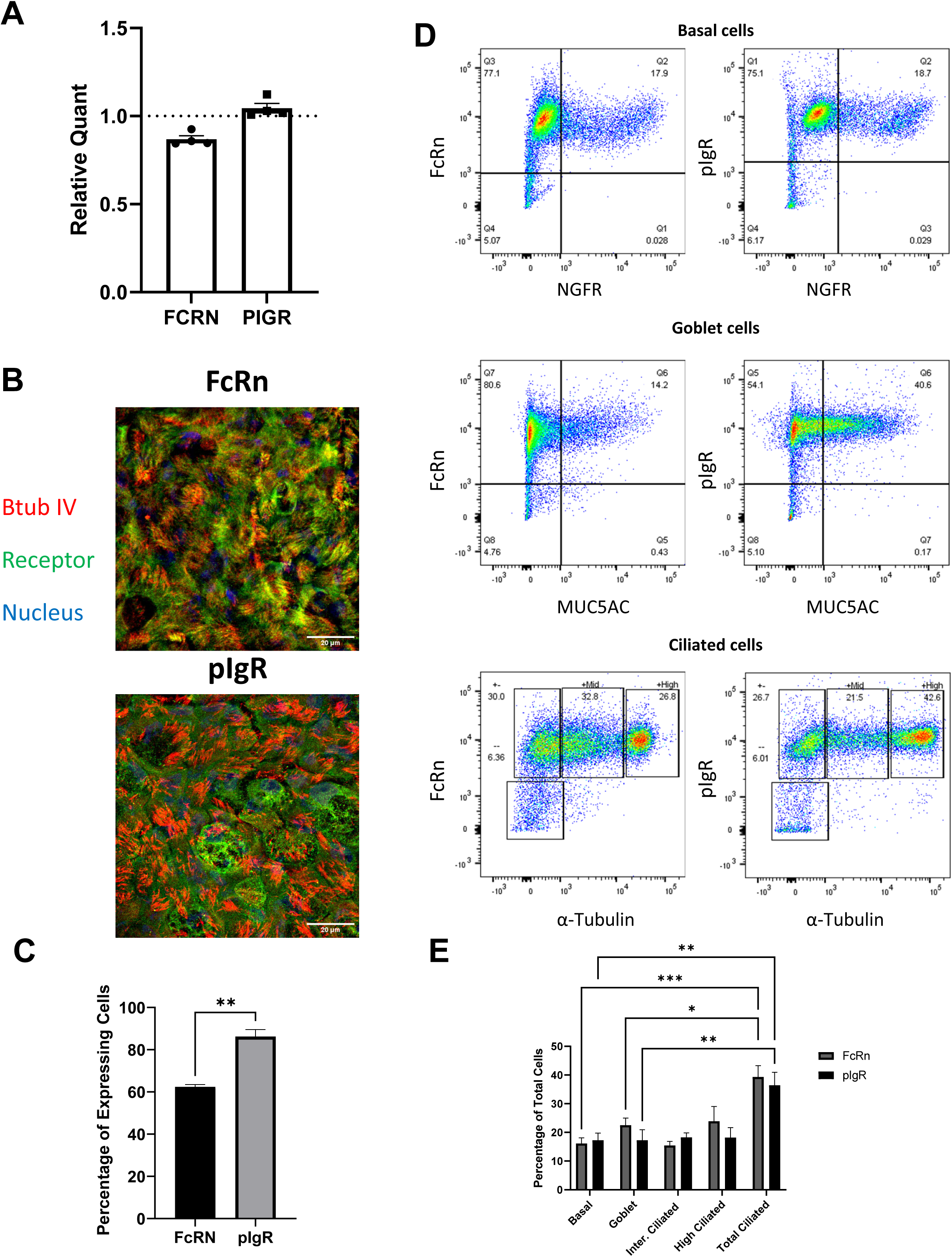
Expression of FcRn and pIgR in human nasal epithelial cells. A) Relative expression levels by qRT-PCR of FcRN and pIgR, normalized to GAPDH. B) Representative images of FcRN and pIgR expressed in hNECs by indirect immunofluorescence microscopy. hNECs were stained for β-tubulin IV (red), FcRn or pIgR (green) and Hoechst (blue). C) Quantification of immunofluorescence images. Total number of cells that expressed FcRn or pIgR in hNECs images (3 images for each with over 300 cells per image). D) Representative flow cytometric density plots of major cell types derived from nasal epithelial cell cultures. Gating schemes for basal cells, Goblet cells and Ciliated cells vs FcRN or pIgR are shown. E) Quantification of flow cytometry cell types by percentage of total cells, values are expressed as mean ± SE, n = 4/group. *P<0.05 **P<0.01 ***P < 0.001 ****P<0.0001.

Quantification of stained cultures demonstrated a significantly greater proportion of cells expressing pIgR compared to FcRn (Figure 1C). Furthermore, to determine the distribution of FcRn and pIgR expression across nasal epithelial cell populations, flow cytometry analysis was performed on differentiated hNECs. Anti-NGFR was used to identify basal cells, anti-MUC5AC for goblet cells, and stained for acetylated α-tubulin for ciliated cells. Both FcRn and pIgR were detected in basal, goblet, intermediate ciliated, and highly ciliated cells (Figure 1D). FcRn expression trended higher in goblet and highly ciliated cells relative to pIgR expression (Figure 1E). In addition, both receptors were highly expressed in ciliated cells compared to basal and goblet cells (Figure 1E). Collectively, these findings confirm expression of both transcytosis receptors in hNEC cultures.

### Purified Antibody Binding Activities to SARS-CoV-2 Proteins

Once the expression of FcRn and pIgR was confirmed in the hNEC cultures, the binding activity of IgG and IgA antibodies purified from convalescent plasma from a patient who was infected with an early variant of SARS-CoV-2, vaccinated, and boosted with the ancestral vaccine was examined to SARS-CoV-2 proteins. To assess purified antibody binding to viral protein, hNECs were infected with a low MOI of SARS-CoV-2 ancestral-D614G, Delta, or Omicron variants and co-immunostained using purified IgG or IgA as a primary antibody and anti-nucleoprotein antibody imaging. Representative images show antibody binding of both IgG and IgA in infected cells but not in uninfected cells (Figure 2A). In addition, purified IgG and IgA were utilized as primary antibodies to stain Vero cells that were transfected with a plasmid encoding the SARS-CoV-2 ancestral D614G spike protein. Representative images showed that transfected cells demonstrated binding efficiency to purified IgG and IgA similarly to infected cells utilizing commercial anti-spike IgG antibodies (Supplemental Figure 2). To further characterize the SARS-CoV-2-specific responses of purified IgG and IgA, a 15-plex MagPix assay was performed to quantify the number of antibodies against spike proteins from multiple variants of concern (VOCs). The results demonstrated that both IgG and IgA antibodies recognized spike proteins from the UK, South African (SA), Delta, and Brazil SARS-CoV-2 VOCs. IgG antibodies exhibited strong binding responses to VOCs at values greater than 1000 U/mL (Figure 2B). In contrast, IgA antibodies displayed anti-spike titers less than 250 U/mL for the SARS-CoV-2 variants (Figure 2B).

**Figure 2.**
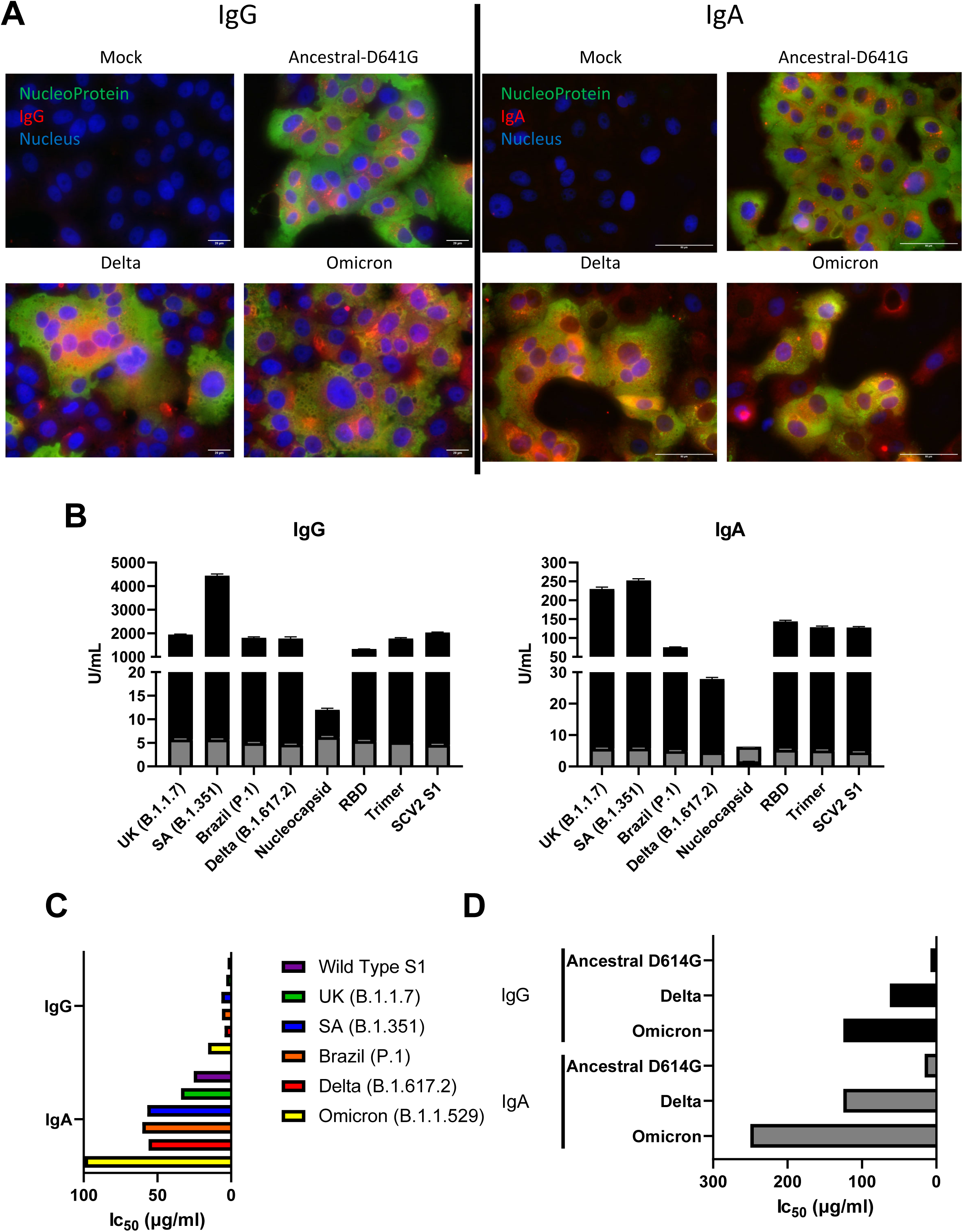
Antibody Response to SARS-CoV-2 Proteins. A) Vero-TMPRSS2 cells were infected with HP07, Delta, or Omicron variants, fixed, and immunostained using purified human IgG or IgA from convalescent plasma. B) 15-plex Magpix assay was conducted on IgG and IgA antibodies that were purified from convalescent plasma. Anti-SARS-CoV-2 protein titers were determined for spike protein of different variants. Antibody levels are expressed as arbitrary units/ml (U/ml). Cutoff lines are represented as gray bars. Values are expressed as mean ± SE, n = 3/Ig. C) IC50 values of ACE2 binding inhibition were conducted using purified IgG or IgA against SARS-CoV-2 spike protein of different variants. D) IC50 values of SARS-CoV-2 neutralization using purified IgG or IgA against SARS-CoV-2 variants.

The assay was used to evaluate antibody binding to nucleocapsid and multiple spike proteins of SARS-CoV-2 variants, including the spike trimer, receptor-binding domain (RBD), and S1. Consistent with previous findings, both IgG and IgA demonstrated binding above the assay cutoff values, with IgG greater than 1000 U/mL and IgA below 250 U/mL (Figure 2B). Furthermore, purified antibodies exhibited cross-reactivity against seasonal human coronavirus spike protein, including CoV-229E, CoV-HKU1, CoV-BL63, CoV-OC43, and SCV S1. IgG antibodies demonstrated cross-reactivity against all seasonal coronaviruses spike proteins (Supplemental Figure 4). In contrast, IgA antibodies do not exhibit detectable cross-reactive binding, with titers remaining below threshold values (Supplemental Figure 4). Overall, purified antibodies from the convalescent plasma of patients who were previously infected and subsequently vaccinated and boosted with SARS-CoV-2 exhibited cross-reactivity to several SARS-CoV-2 proteins.

To further evaluate antibody functionality, 6-Plex Magpix assay was performed to determine the ability of purified IgG and IgA to inhibit ACE-2 binding by SARS-CoV-2 spike proteins from multiple variants. The variants examined included ancestral, and Alpha, Beta, Gamma, Delta, and Omicron variants. Overall, purified IgG efficiently inhibited ACE-2 binding of spike protein at IC50 values ranging from 2-15 U/mL, with significant inhibitory activity against Omicron spike protein (Figure 2C). In comparison, IgA inhibited ACE2 binding less effectively with IC50 values of 25-100 U/ml (Figure 2C). To further assess neutralizing activity, purified antibodies were evaluated using microneutralization assays. Serially diluted antibodies were co-incubated with Ancestral D641G, Delta, and Omicron variants for 1 hr prior to infection of Vero/TMPRSS2 cells. Purified IgG had IC_50_ values of 7.8 µg/ml, 62.5 µg/ml, and 125 µg/ml for Ancestral D641G, Delta, and Omicron variants, respectively (Figure 2D). Compared with IgG, IgA IC_50_ values were less efficient at neutralizing SARS-CoV-2 with IC50 values of 15.6 µg/ml, 125 µg/ml, and 250 µg/ml for Ancestral D641G, Delta, and Omicron variants, respectively (Figure 2D). Together, these findings characterize the neutralization capacity of purified IgG and IgA against multiple SARS-CoV-2 VOCs.

### Antibody Transcytosis in Human Nasal Epithelial Cells

Transepithelial electrical resistance measurements were obtained for each well to determine cell barrier integrity. TEER results showed that hNECs exhibited resistance values greater than the minimum resistance for positive cell barrier integrity, 500 ohms (Figure 3A). To ensure that transcytosis of IgG and IgA is occurring by a receptor-mediated transport rather than passive leakage across the epithelial barrier, permeability of the hNECs cultures was assessed using the transepithelial passage of a fluorescent tracer FITC-Dextran. FITC-Dextran was administered to the apical compartment of the trans-wells, and the fluorescence intensity of the basal compartment was measured using a spectrophotometer. The data showed a significant decrease in relative fluorescence intensity in differentiated hNECs cultures compared to empty Trans-well controls, further confirming that the epithelial barrier of the hNECs cultures remained intact (Figure 3B).

**Figure 3.**
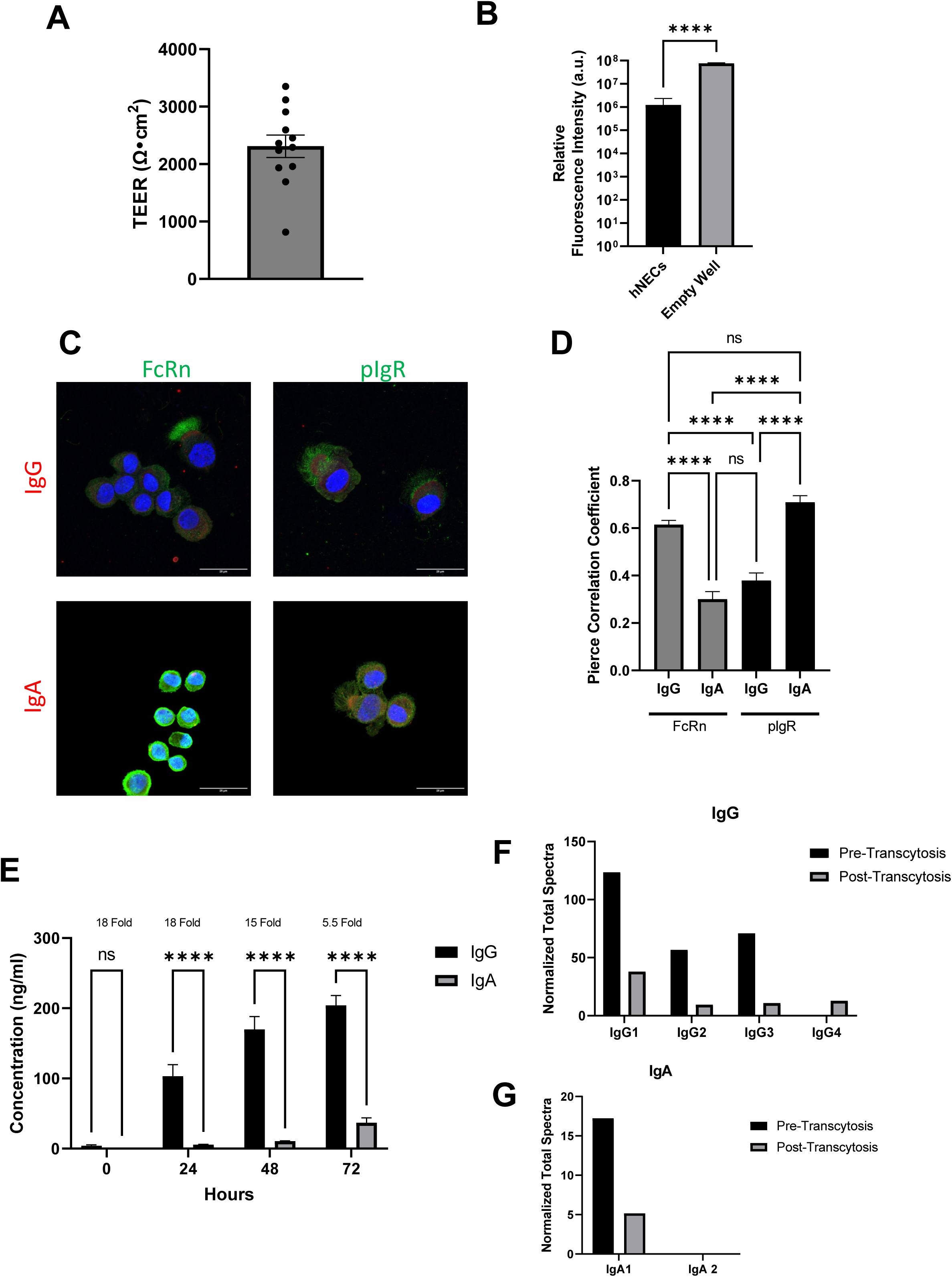
Antibody transcytosis in human nasal epithelial cells. A) Trans Epithelial Electrical Resistance (TEER) measurements were conducted on fully differentiated hNECs. Values are expressed as mean ± SE, n = 12 wells. B) FITC-dextran permeability in hNECs was assessed in basolateral treated compartments. Values are normalized against samples that were placed on the apical side and expressed as mean ± SE, n = 12 wells. C) Representative images of hNECs lifted, smeared on a slide and immunostained for IgG/IgA and FcRn/pIgR. D) Pierce Colocalization analysis was conducted on these images. Values are expressed as Pierce Coefficients, n= 10. E) Transcytosis of IgG or IgA was determined using an ELISA assay. Values are expressed as mean ± SE, n = 4well/time point. ****P<0.0001. Breakdown of IgG subclasses (F) or IgA (G) in the starting sample (pre-transcytosis) or transcytosed to the apical surface of hNECs.

In polarized epithelial cells, human FcRn mediates bidirectional transcytosis of IgG, whereas pIgR facilitates unidirectional transport of polymeric IgA and IgM ^(23,91)^. To gain insight into the ability of these receptors to bind and transport antibodies in hNEC cultures, confocal microscopy colocalization experiments were performed. Purified IgG or IgA was added to the basolateral compartment of differentiated hNECs cultures. After 48 hr, hNECs were trypsinized and spun onto slides. Immunofluorescence co-staining was conducted using anti-human IgG or IgA and anti- FcRn or pIgR antibodies (Figure 3C and 3D). For FcRn, the Pierce correlation coefficient was higher with IgG compared to IgA (Figure 3D). Conversely, pIgR demonstrated a higher Pierc correlation with IgA, compared to IgG (Figure 3D). To further validate these findings, similar experiments were conducted in non-trypsinized hNECs cultures. Consistent with the single-cell analysis, FcRn:IgG and pIgR:IgA colocalized, whereas reciprocal receptor-antibody complexes exhibited minimal colocalization (Supplemental Figure 5).

Subsequently, to quantify antibody transcytosis across hNECs, purified IgG or IgA antibodies were added to the basolateral compartment, and apical washes were collected every 24 h for 72 h. Anti-human IgG and IgA ELISAs were performed to determine the concentration of transcytosed antibodies. Concentrations of transcytosed IgG detected in the apical compartment were significantly greater than those of IgA at 24, 48, and 72 h (Figure 3E).

Although IgG transport remained greater throughout the experiment, the fold difference between IgG and IgA transcytosis decreased by 72 h (Figure 3E). The transcytosed IgG and IgA wer quantified for their subclass and the amount of IgG subclasses transcytosed were reflective of the amount present in the starting pool (Figure 3F). Only IgA1 was detected in the starting pool of purified IgA and so that was transcytosed exclusively (Figure 3G). Taken together, these findings indicate that antibodies are transported from the basolateral to the apical compartment in differentiated hNEC cultures.

### Transcytosed Antibodies Neutralize Viral Infection in Human Nasal Epithelial Cells

Having established that the hNEC cultures are capable of transcytosing purified antibodies, the ability of transcytosed antibodies to retain antibody activity and neutralize viral infection was next evaluated. Differentiated hNECs were treated basolaterally for 24h with increasing concentrations of purified IgG or IgA isolated from convalescent or pre-pandemic plasma before SARS-CoV-2 infection.

Apical washes were collected at the indicated time points to quantify infectious virus production by TCID_50_/mL. IgG concentrations greater than 40 µg/ml and IgA concentrations greater than 200 µg/ml significantly reduced viral titers compared to untreated controls. (Figure 4A). To assess neutralization kinetics over time, hNECs were treated basolaterally with 1000 µg/ml purified IgG or IgA before apical SARS-CoV-2 infection. Both IgG- and IgA-treated cultures exhibited significantly reduced viral titers at all examined time points compared to untreated controls, with titers approaching the limit of detection (Figure 4B). In contrast, antibodies purified from pre-pandemic plasma did not significantly reduce viral titers relative to untreated controls (Figure 4B).

**Figure 4.**
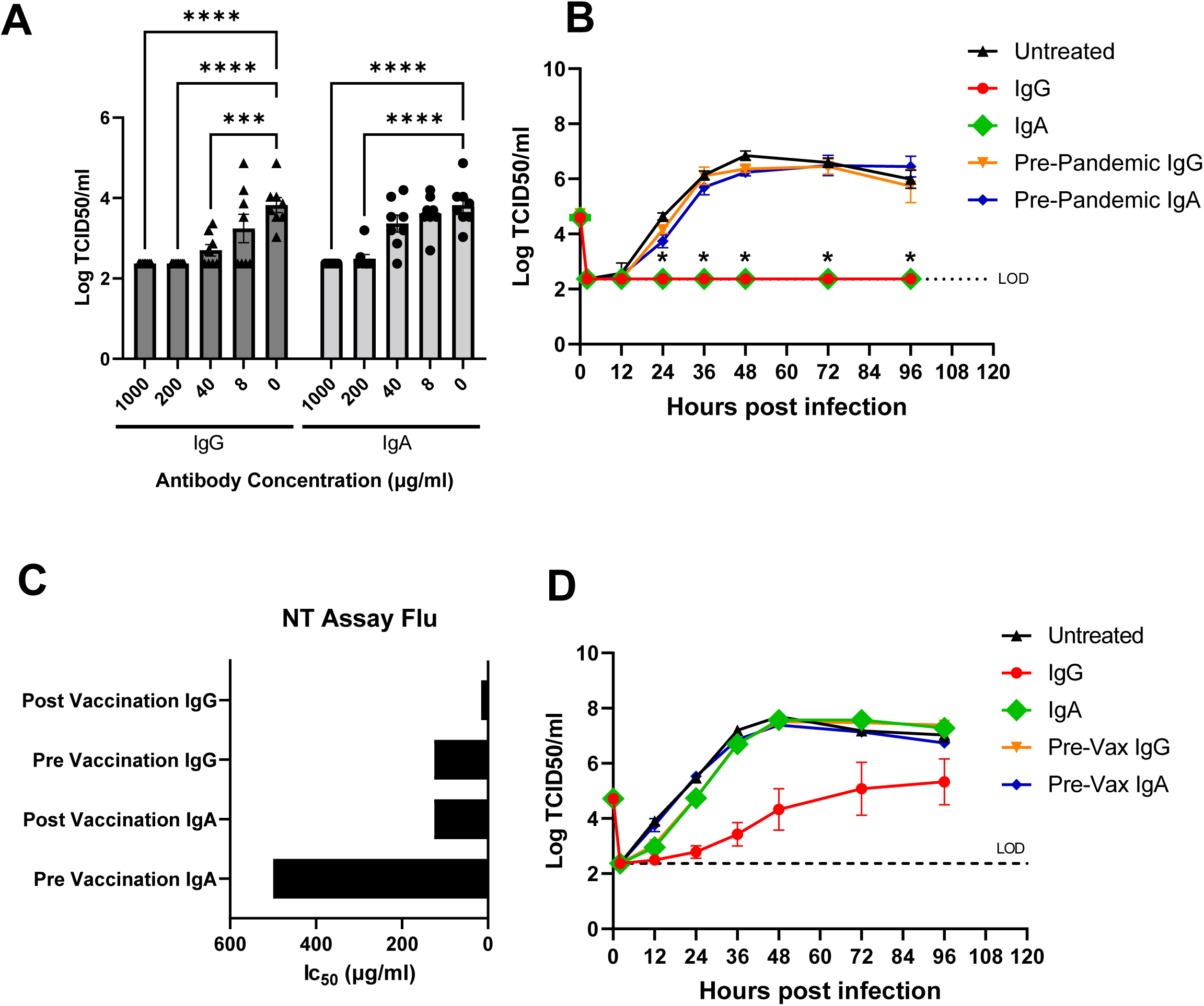
Neutralization of SARS-CoV2 and Influenza A Virus by transcytosed IgG or IgA. A) 24-hour post infection infectious virus titers were obtained from apical washings of hNECs infected with SARS-CoV-2 ancestral-D614G variant and pretreated with different concentrations of IgG or IgA in the basolateral compartment 24 hours before infection. Data are pooled from three independent experiments with n=8 wells per concentration (total n=40 wells per antibody type). B) Low MOI, multistep growth curves were performed on hNEC cultures with 1000 ug/ml of purified IgG or IgA placed on the basolateral compartment added 24 hours before infection and infected with SARS-CoV-2 (Ancestral-D641G). Data pooled from n=8 wells per antibody. C) IC50 values of influenza inhibition were collected using purified IgG or IgA against vaccine H3N2 influenza virus. D) Low MOI, multistep growth curves were performed on hNEC cultures with 1000 ug/ml of purified IgG or IgA placed on the basolateral compartment 24 hour before infection and infected with vaccine H3N2 influenza virus. Data pooled from n=8 wells per antibody. ***P < 0.001 ****P<0.0001.

Given the neutralization observed against SARS-CoV-2, influenza virus neutralization was next evaluated using antibodies purified from plasma collected before or after influenza vaccination. Serial dilutions of purified antibodies were co-incubated with the influenza virus before infection of MDCK-SIAT cells. Post-vaccination IgG exhibited greater neutralization than pre-vaccination IgG, with IC_50_ values of 15.6 µg/ml and 125 µg/ml, respectively (Figure 4C). In comparison, IgA demonstrated lower neutralization activity, with pre- and post-vaccination IC_50_ values of 500 µg/mL and 125 µg/mL, respectively (Figure 4C). The ability of IgG and IgA to transcytose across differentiated hNECs and neutralize influenza infection over time was then evaluated.Basolateral compartments of hNECs were treated for 24 h with 1000 µg/mL purified IgG or IgA isolated from pre- or post-vaccination purified IgG or IgA before infection with influenza A/H3N2 (A/Kansas/X-327). Apical washes were collected at the indicated time points following infection to quantify viral titers. Treatment with post-vaccination IgG resulted in significantly reduced viral titers beginning at 24 h post-infection compared to untreated controls and pre-vaccination IgG-treated cultures (Figure 4D). In contrast, IgA purified from vaccinated individuals did not significantly reduce viral titers compared to untreated or pre-vaccination IgA-treated cultures (Figure 4D).

### Distinct FcRn and pIgR Expression Profiles in Upper and Lower Respiratory Tract

Previous studies have demonstrated that the upper and lower respiratory tract possess distinct physiological functions, cellular compositions, and immune responses that reflect their specialized roles within the respiratory system ^(93–95)^. Although the upper respiratory tract serves as the initial site of infection for many respiratory pathogens, infection frequently progresses to the lower respiratory tract. To investigate the ability of the lower respiratory tract epithelium to transcytose antibodies, primary differentiated human bronchial epithelial cells (hBECs) were utilized. Expression of FcRn and pIgR proteins in differentiated hBECs was confirmed by Western blot analysis (Figure 5A). Compared to hNECs, hBECs exhibited significantly increased FcRn protein expression and reduced pIgR protein expression (Figure 5B, 5C).

**Figure 5.**
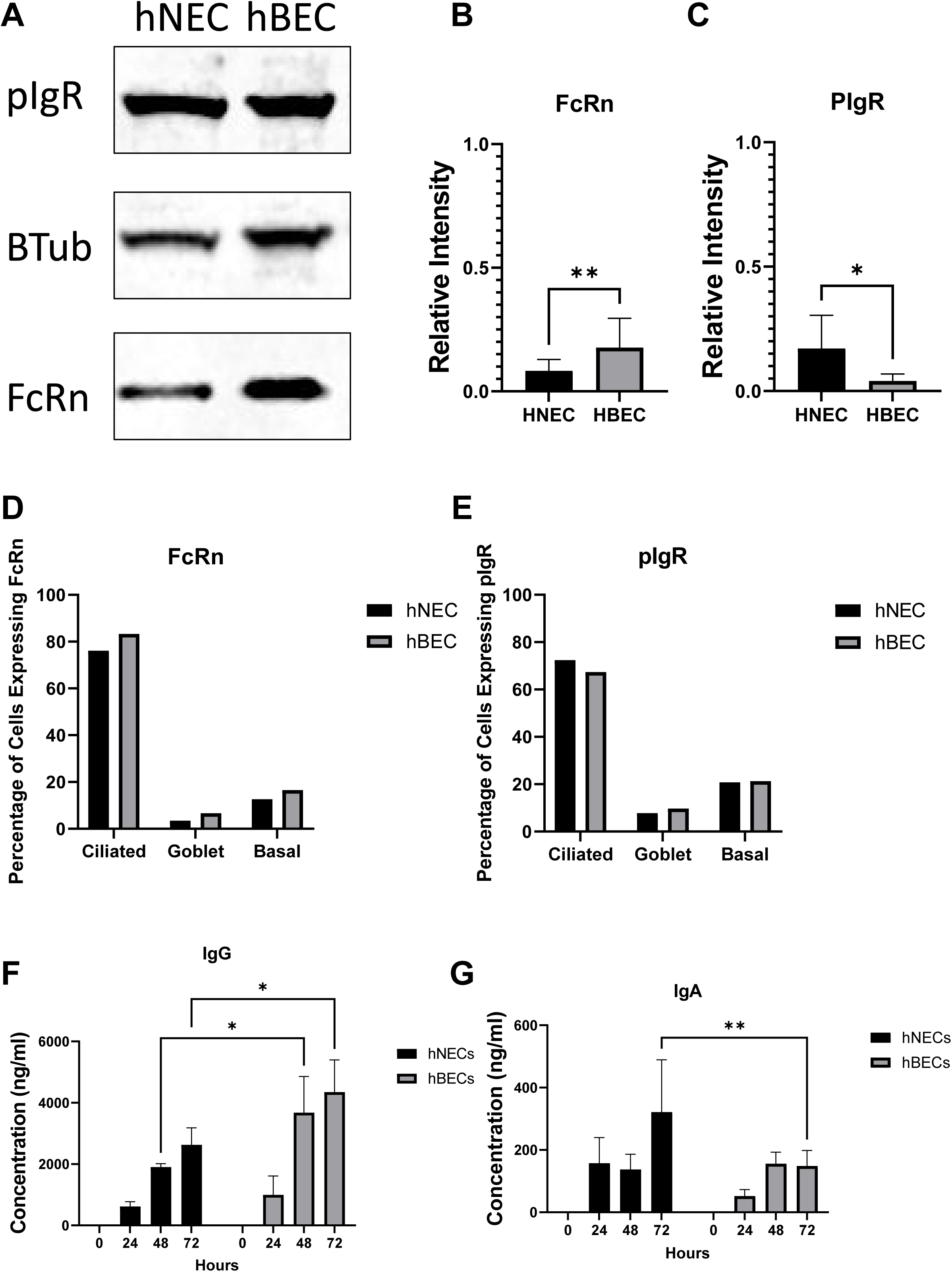
Human nasal and bronchial epithelial cells exhibit the ability to transcytose IgG and IgA. A) Immunoblot of FcRn and pIgR in hNECs and hBECs. B, C) Relative protein expression levels by Western blotting of FcRN, pIgR, normalized to β-tublin, n = <3. D, E) Quantification of flow cytometry cell types by percentage of total cells, values are expressed as mean ± SE, n = 1/group. F, G) Transcytosis of IgG and IgA was determined using an ELISA assay. Values are expressed as mean ± SE, n = <3well/time point. *P<0.05 **P<0.01

To determine whether FcRn and pIgR expression across epithelial cell populations in hBECs was comparable to that observed in hNECs, flow cytometry analyses were performed in differentiated hBECs cultures. Similar patterns of FcRn and pIgR expression were observed across ciliated, goblet, and basal cell populations in both epithelial culture systems (Figure 5D, 5E). Interestingly, although the proportions of cells expressing FcRn and pIgR were comparable between hBECs and hNECs, differences were observed in antibody transcytosis efficiency between the two epithelial models. As demonstrated above in hNECs, both purified IgG and IgA antibodies were capable of transcytosing from the basolateral to the apical compartment.

However, in hBECs cultures, significantly greater concentrations of transcytosed IgG were detected compared to IgA at 48 hr and 72 hr post-treatment (Figure 5F). Furthermore, hNECs exhibited greater IgA transcytosis compared to hBECs at 72 hr, suggesting regional differences in antibody transport across upper and lower respiratory epithelial cells (Figure 5G).

## Discussion

Intramuscular vaccination induces systemic neutralizing antibodies against respiratory viruses but the mechanisms by which these antibodies access the airway lumen and contribute to mucosal protection remain incompletely understood. In this study, FcRn- and pIgR-mediated antibody transcytosis was demonstrated in differentiated human respiratory epithelial cultures, supporting a role for epithelial antibody transport in antiviral mucosal immunity, particularly during SARS-COV-2 infection.

Expression of human FcRn and pIgR was detected in primary differentiated human nasal and bronchial epithelial cell cultures. FcRn and pIgR are expressed on the mucosal surfaces of both the airway and intestinal epithelium ^(42–49)^. Additional characterization of receptor distribution across epithelial cell populations within the nasal and bronchial epithelium demonstrated a greater proportion of ciliated cells expressed FcRn and pIgR relative to goblet and basal cell populations. Similar localizations of these receptors along the apical surface of polarized respiratory epithelium have been described previously ^(48–52)^.

Anti-spike antibodies play a critical role in SARS-CoV-2 clearance, as high levels of neutralization antibodies induced by vaccination have been identified as strong correlates of protection against infection ^(40,96)^. Purified IgG and IgA antibodies isolated from convalescent plasma demonstrated specificity for the SARS-CoV-2 spike protein. However, purified IgG preparations contained higher anti-spike antibody titers compared to purified IgA preparations. These findings are consistent with previous studies demonstrating that the majority of SARS-CoV-2 spike-specific antibody responses are predominantly of the IgG isotype ^(53–56)^. In addition, serum IgA responses have been reported to decline more rapidly following SARS-CoV-2 vaccination or infection and to exhibit lower neutralization potency compared to IgG antibodies ^(57,58)^. Consistent with these observations, ACE2 inhibition assays further corroborated these findings, with lower IC_50_ values for IgG antibodies relative to IgA antibodies, indicating greater inhibitory activity against SARS-CoV-2 spike-ACE2 interactions.

Microneutralization assays are typically performed under in vitro conditions and may not fully recapitulate viral neutralization within the respiratory tract environment. The establishment of hNEC and hBEC cultures, together with the demonstration of antibody transcytosis activity, provides a physiologically relevant model for studying the unique epithelial environment during respiratory viral infection. Differential expressions of transcytosis receptors were observed between epithelial culture systems, with hNECs exhibiting greater pIgR expression relative to FcRn, whereas hBECs demonstrated increased FcRn expression relative to pIgR. In both epithelial culture systems, ciliated cell populations exhibited the highest expression of FcRn and pIgR compared to goblet and basal cell populations. These findings differ from previous studies utilizing explanted lungs from patients with chronic obstructive pulmonary disease (COPD) and donor lungs declined for transplantation, in which single-cell RNA sequencing (scRNA-seq) and immunostaining analyses identified secretory cells as the predominant source of pIgR expression within the small airway epithelium ^(97)^. Differences between these findings may be attributable to the distinct anatomical regions examined, as the previous study focused primarily on distal small airways, methodological differences, such as scRNA-seq and RNA-ISH analysis of explanted tissues.

Although both epithelial culture systems demonstrated antibody transcytosis activity, the hBECs exhibited greater IgG transcytosis compared to hNECs. In contrast, transcytosed IgA concentrations were greater in hNECs compared to hBECs at later time points. In normal human serum, approximately 80% of total antibodies are IgG, whereas IgA comprises roughly 15% of circulating immunoglobulins. Moreover, nearly 90% of serum IgA exists in monomeric form, while only approximately 10% is present as dimeric IgA ^(66)^. Because pIgR preferentially binds to dimeric IgA, this would imply that the smaller concentration of transcytosed IgA could be a result of the significantly smaller proportion of dimeric IgA found in plasma. In addition, functional differences between serum-derived IgA and mucosal secretory IgA, including differences in receptor interactions, may also contribute to differences in IgA transcytosis efficiency. Lastly, it is important to note that FcRn-mediated IgG transport occurs bidirectionally, whereas pIgR-mediated IgA transport is primarily unidirectional toward the lumen, which may ultimately promote greater accumulation of IgA at mucosal surfaces over time. Additional studies are required to further define the role of FcRn- and pIgR-mediated transport directionality in regulating antibody distribution within the nasal and bronchial epithelium and antibody concentrations in the airway lumen.

Colocalization of IgG and IgA antibodies against FcRn and pIgR, respectively, provides additional support for a receptor-mediated antibody transport within these primary epithelial culture systems. It is important to acknowledge that FcRn- and pIgR-mediated transport is sensitive to the pH of the environment for strong binding; therefore, future studies will be required to further evaluate the role of pH within primary epithelial cultures during antibody transcytosis and early viral infection.

Given that SARS-CoV-2 primarily infects the upper respiratory tract, mucosal immunity has a major role in viral clearance ^(77)^. Mucosal antibodies represent one of the primary lines of defense against viral infection at these mucosal surfaces ^(78,79)^. At the mucosal interfaces, antibodies can limit infection by blocking viral attachment, entry, and replication within epithelial cells. In the present study, we hypothesized that antibodies introduced into the basolateral compartment of our hNECs cultures would undergo transcytosis to the apical side, where they could mediate neutralization of SARS-CoV-2 infection. The results demonstrated that both purified IgG and IgA were capable of transcytosing and neutralizing SARS-CoV-2 infection. IgG exhibited a greater potency to neutralize SARS-CoV-2 than IgA, with significant neutralization observed at concentrations of 40 µg/mL, whereas IgA-mediated neutralization required concentrations up to 200 µg/mL. The differences in neutralization efficiency may reflect the differences in the anti-spike IgG vs IgA antibody titers present within the convalescent plasma preparations. In addition, the concentration of dimeric IgA that is transcytosed is significantly less than that of IgG; therefore, there is less antibody available to neutralize the virus. Despite these differences, treatment with 1000 µg/ml of purified antibodies resulted in no difference in SARS-CoV-2 titers for both IgG- and IgA-treated hNECs. SARS-CoV-2 infection elicits virus-specific IgG and IgA responses within saliva and bronchoalveolar fluid that differ from systemic plasma antibody responses. Furthermore, these differences may also reflect distinct B-cell populations responsible for antibody production, such as tissue-resident memory B cells at barrier tissue surfaces and memory B-cell populations within secondary lymphoid organs. As the primary form of IgA within the nasopharynx, dimeric IgA has been shown to have an enhanced ability to neutralize SARS-CoV-2 compared to IgG ^(80,81)^. Therefore, further studies on investigating vaccination strategies capable of inducing a stronger mucosal antibody repertoire and durable immunity at the respiratory surfaces are warranted.

Transcytosed IgG antibodies mediated partial neutralization of influenza A infection in hNECs. In contrast, IgA antibodies did not produce a significant reduction in viral titers compared to untreated or unvaccinated controls ^(82–84)^. It is important to note that viral titers of influenza A are higher than those of SARS-CoV-2 in untreated controls, which may additionally contribute to the reduced neutralization observed. These results point towards virus-specific factors that may influence the effectiveness of antibody-mediated neutralization following epithelial transcytosis. An additional factor that may contribute to mucosal immunity is the ability of antibodies to neutralize viruses intracellularly within epithelial cells, which warrants further studies. During transcytosis, dimeric IgA and pentameric IgM have been shown to mediate intracellular neutralization of pathogens, including rotavirus, influenza virus, and HIV, followed by transport of immune complexes back into the lumen, thereby limiting cytolytic damage to the epithelium ^(85–89)^.

In summary, our findings indicate that viral neutralization in the nasal epithelium is influenced by antibody specificity, transcytosis efficiency, and viral replication kinetics. These findings demonstrate that FcRn- and pIgR-mediated antibody transcytosis within the respiratory epithelium contributes to mucosal antiviral immunity against both SARS-CoV-2 and IAV. The efficiency of antibody-mediated neutralization following transcytosis appears to be influenced by multiple factors, including antibody isotype composition, receptor-mediated transport efficiency, viral replication kinetics, and virus-specific susceptibility to neutralization. Furthermore, the establishment of differentiated human respiratory epithelial culture systems provides a physiologically relevant platform for investigating mucosal antibody transport and antiviral activity at barrier surfaces.

## Acknowledgements

The work was supported by National Institutes of Health, Naitional Institute of Allergy and Infectious Disease contract 75N93021C00045 (AP) and T32 AI007417 (NS), a contract (W911QY2090012, DS) with the Joint Program Executive Office for Chemical, Biological, Radiological, and Nuclear Defense of the Department of Defense, Bloomberg Philanthropies, the State of Maryland and The Richard Eliasberg Family Foundation. We thank all the members of the Pekosz laboratory for discussions regarding the data.

