## supplemental figures for "Antibody Transcytosis and Neutralizing Activity in Respiratory Epithelial Cells"

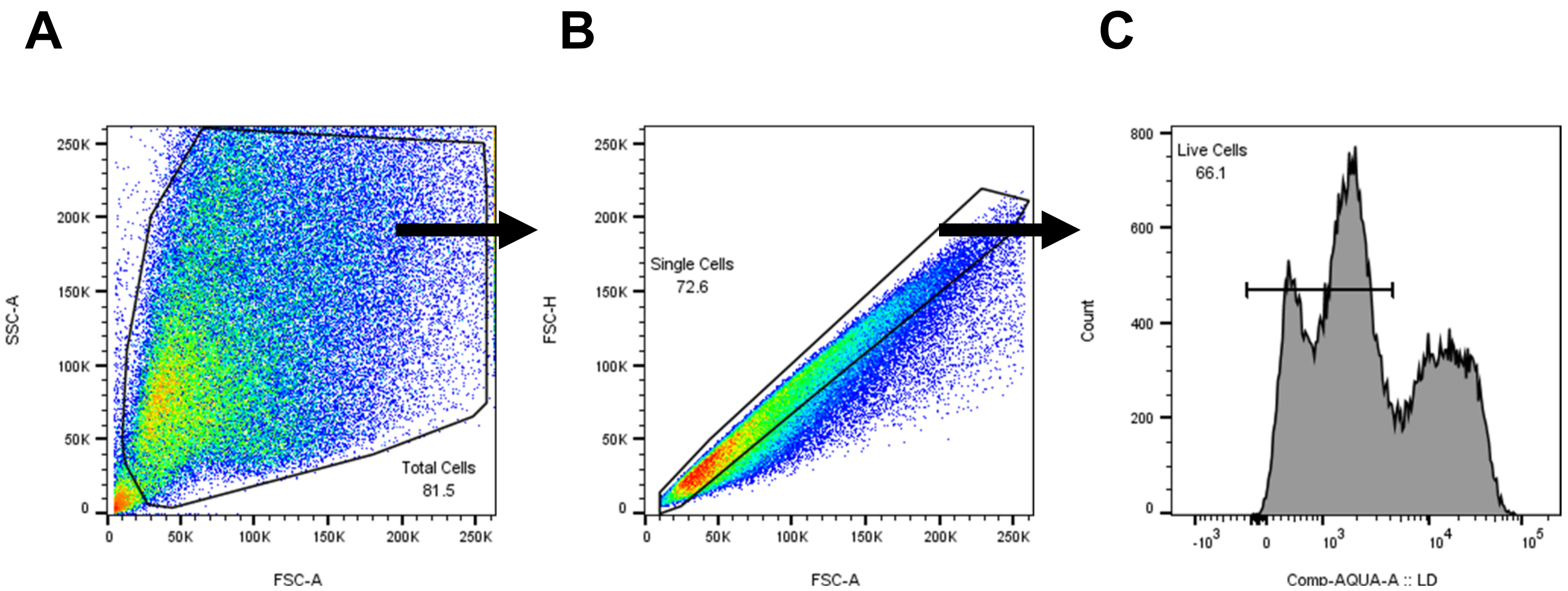


**Supplemental Figure 1**

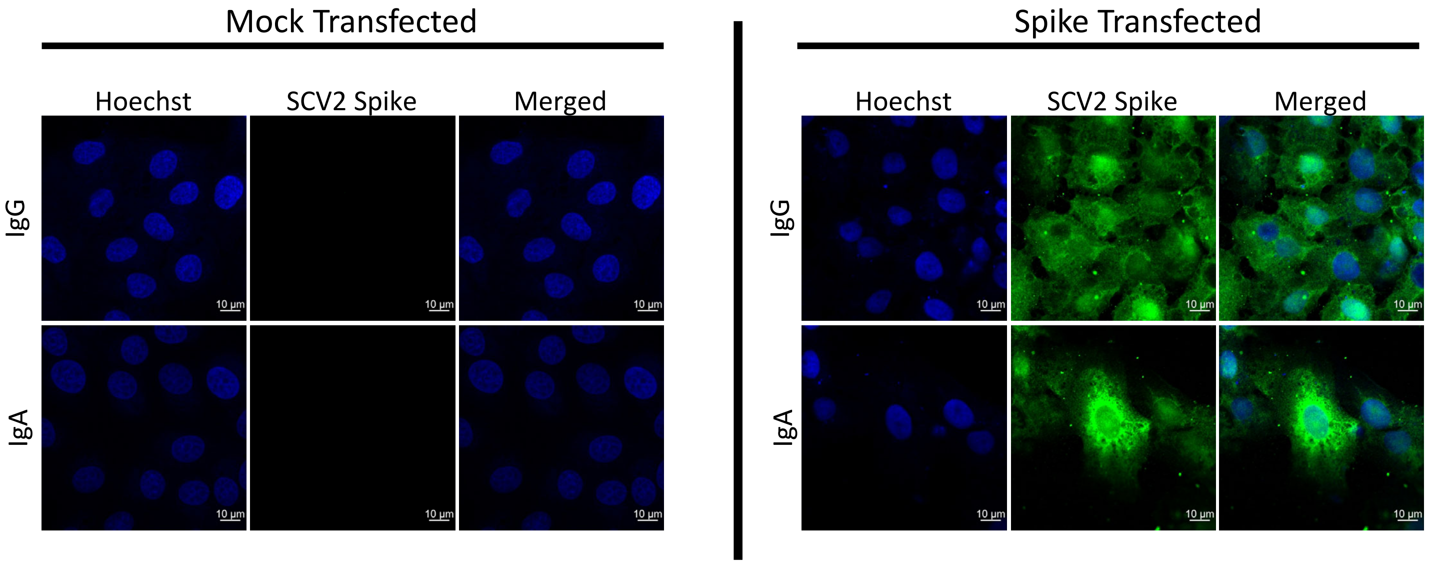


**Supplemental Figure 2.**

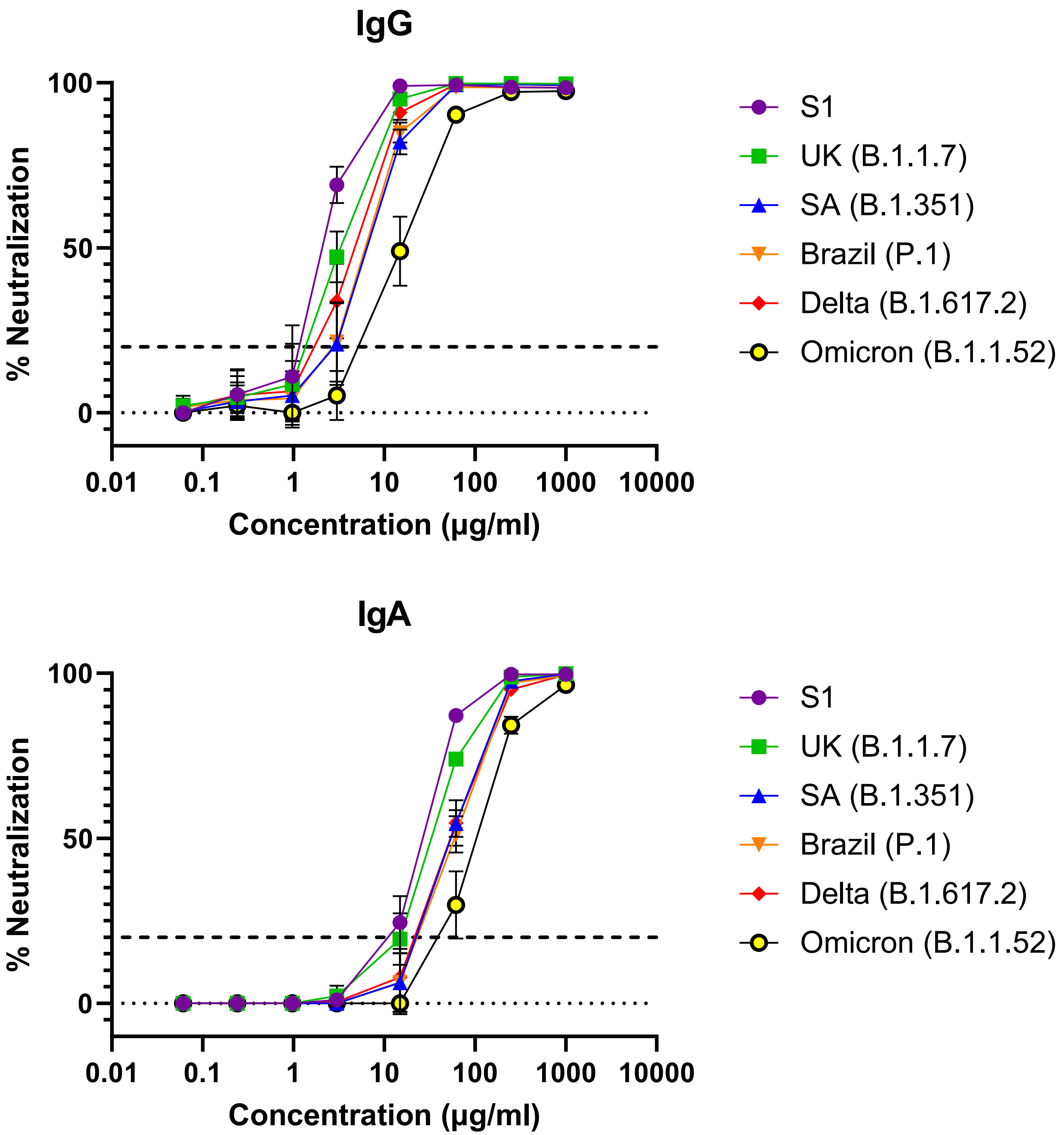


**Supplemental Figure 3.**

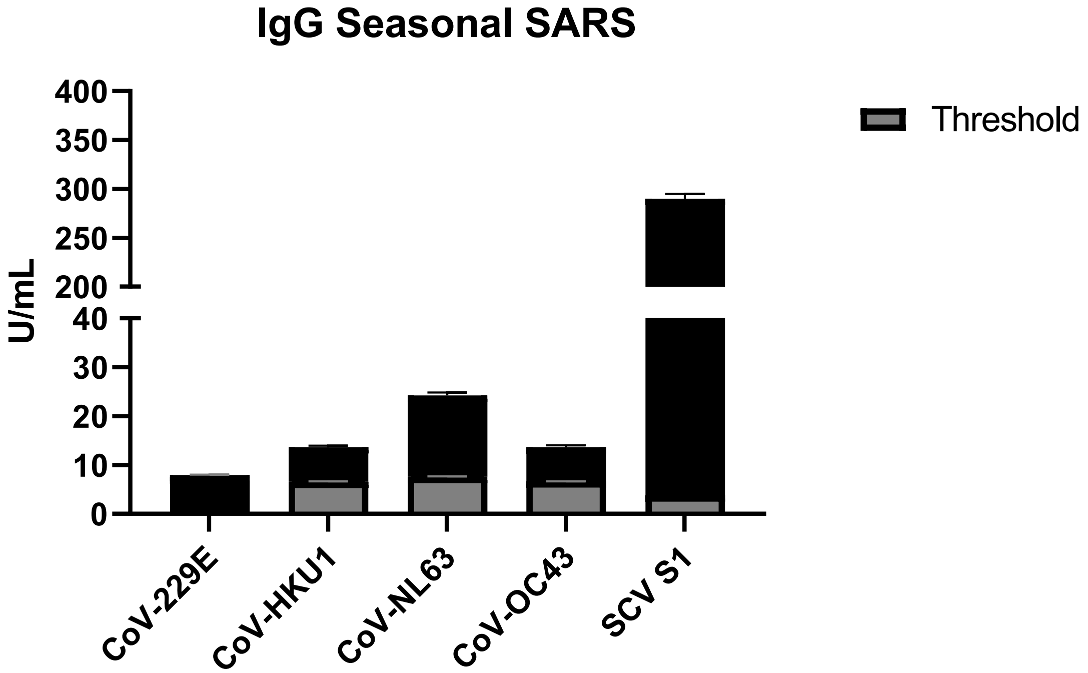


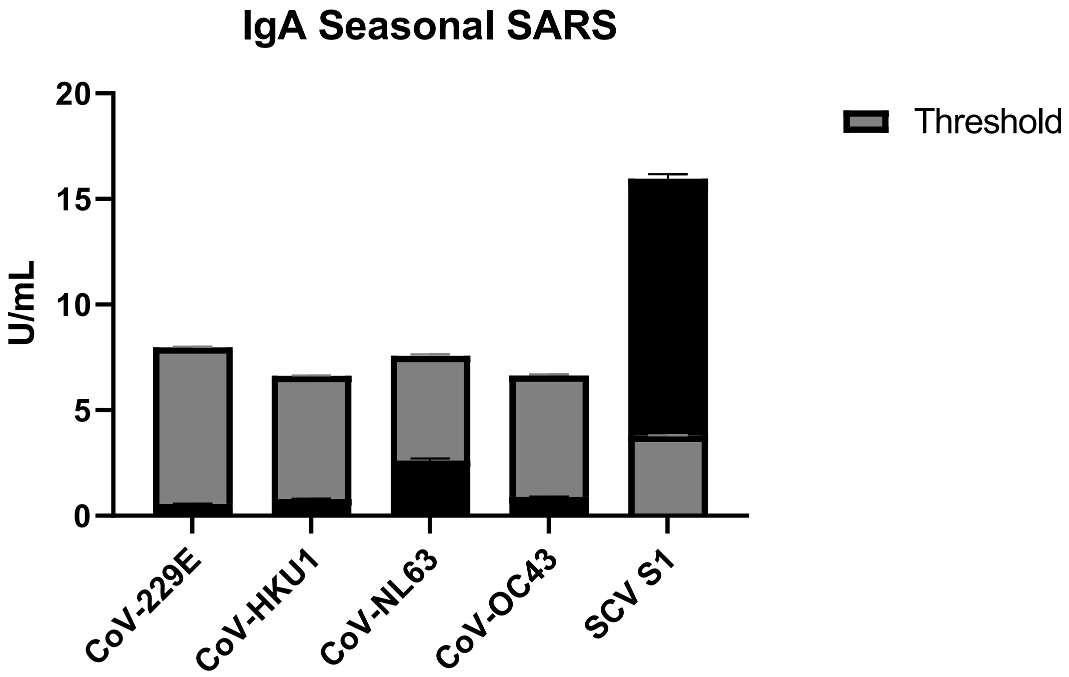


**Supplemental Figure 4**

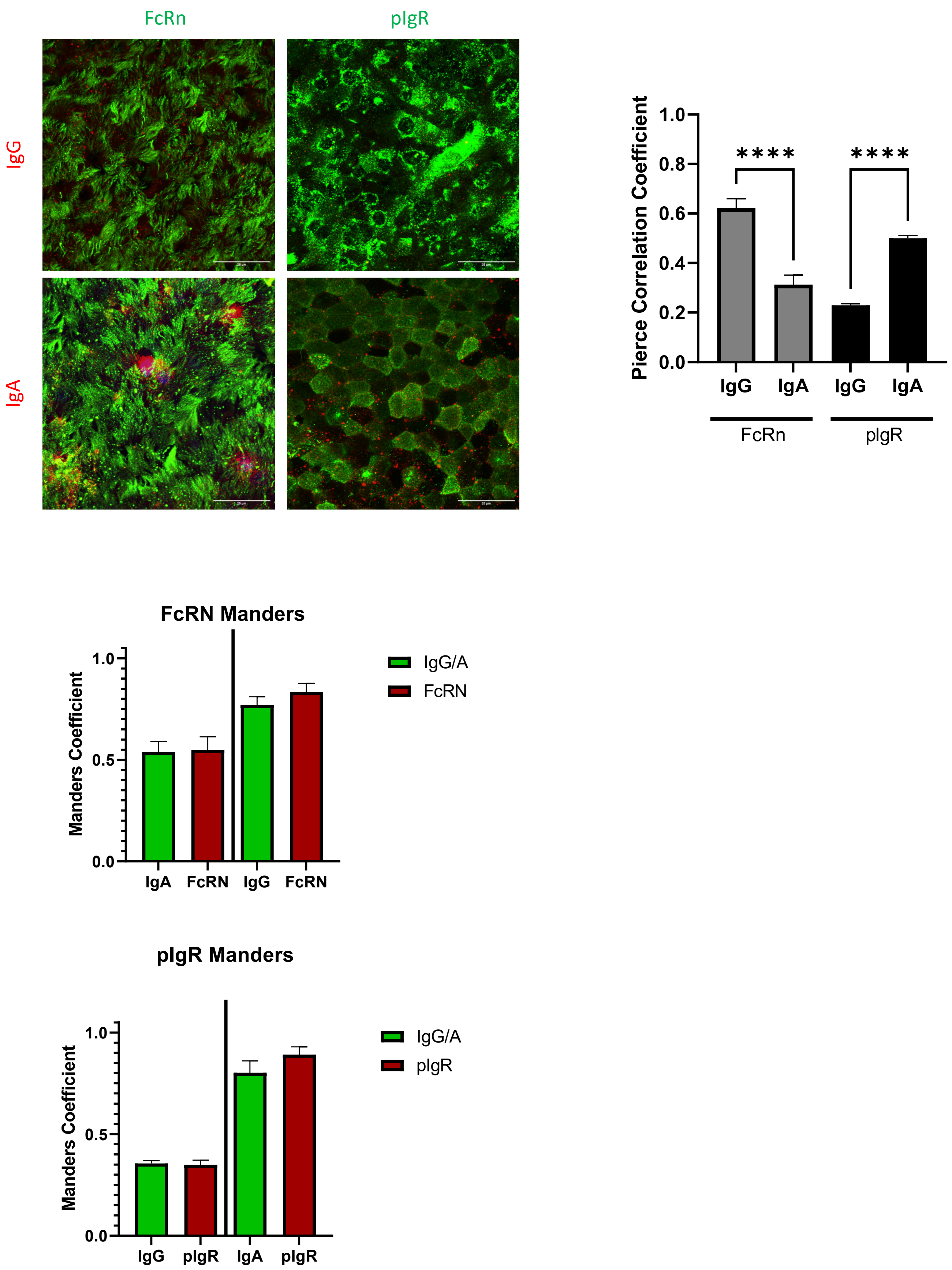


**Supplemental Figure 5.**

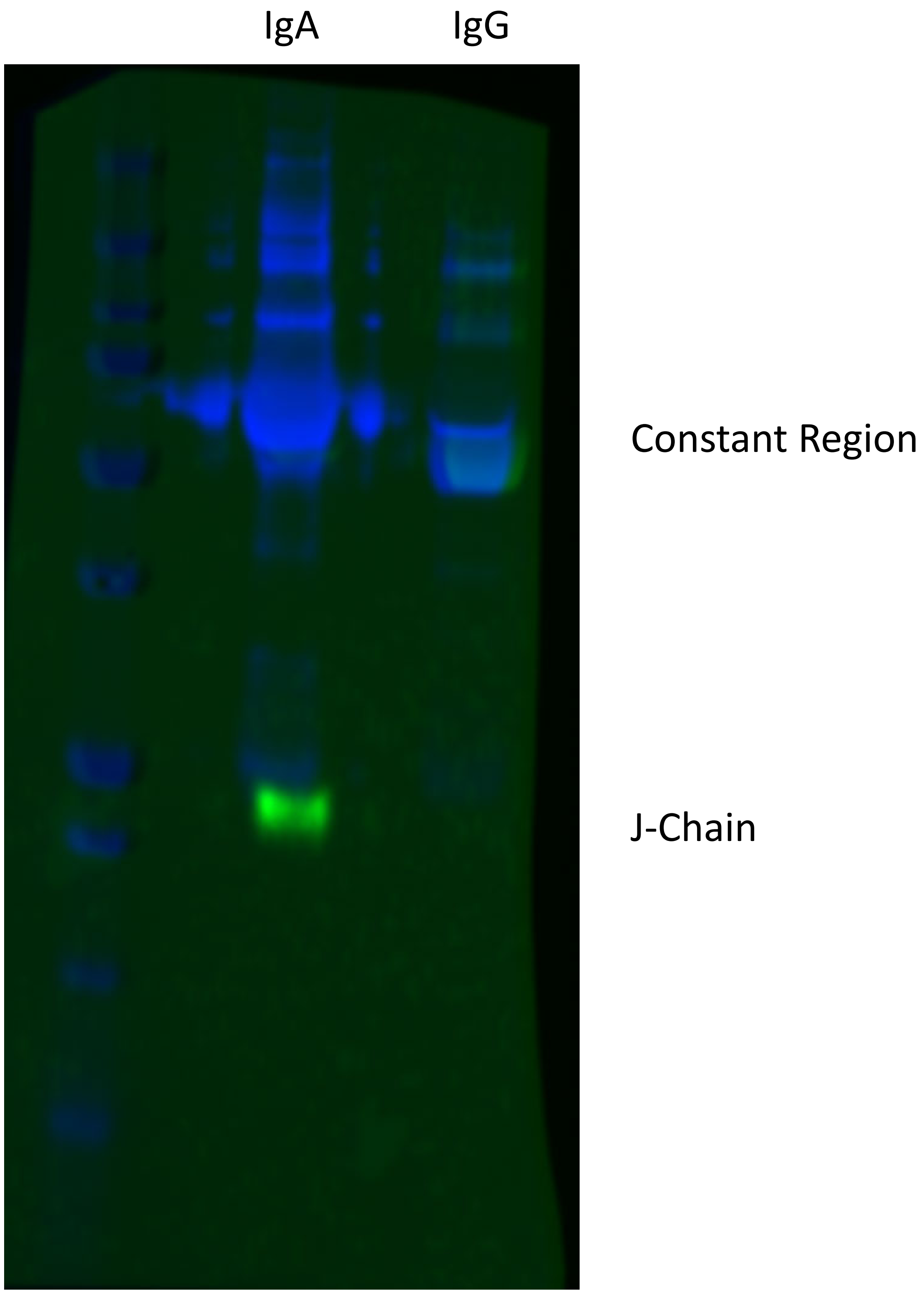


**Supplemental Figure 6.**
